# Mitochondrial decline in the ageing old world primate retina: little evidence for difference between the centre and periphery

**DOI:** 10.1101/2022.08.17.504343

**Authors:** Jaimie Hoh Kam, Harpreet Shinhmar, Michael Barry Powner, Matthew JohnHayes, Asmaa Aboelnour, Glen Jeffery

## Abstract

Mitochondrial decline is a key feature of ageing. The retina has more mitochondria than any other tissue and ages rapidly. To understand human retinal ageing it is critical to examine old world primates that have similar visual systems to humans, and do so across central and peripheral regions, as there is evidence for early central decline. Hence, we examine mitochondrial metrics in young and ageing *Macaca fascicularis* retinae. In spite of reduced ATP with age, primate mitochondrial complex activity did not decline. But mitochondrial membrane potentials were reduced significantly, and concomitantly, mitochondrial membrane permeability increased. The mitochondrial marker Tom20 declined significantly, consistent with reduced mitochondria number, while VDAC, a voltage dependent anion channel and diffusion pore associated with apoptosis increased significantly. In spite of these clear age-related changes, there was almost no evidence for regional differences between the centre and the periphery.

Primate cones do not die with age, but many showed marked structural decline with vacuous spaces in proximal inner segments normally occupied by endoplasmic reticulum (ER), that regulate mitochondrial autophagy. In many peripheral cones, ER was displaced by the nucleus that transposed across the outer limiting membrane and could become embedded in mitochondrial populations. These data are consistent with significant changes in retinal mitochondria in old world primate ageing but provide little if any evidence that aged central mitochondria suffer more than those in the periphery.

## Introduction

Mitochondria regulate metabolism and ageing (1,2). The significance of these features come together in the outer retina that has the highest metabolic rate in the body, the greatest concentration of mitochondria in any tissue and also a very rapid rate of ageing (3,4). Further, as vision is the primary sensory modality for primates, mitochondrial function in the retina underpins our lives. However, the rapid pace of retinal ageing is problematic in relation to progressive extensions to human lifespan. For much of our evolution, lifespan was probably under 40 years, but is now more than double that in the Western world and parts of Asia. This extension exposes the vulnerability of the retina (5,6).

By the age of approximately 70 years, around 30% of human central rod photoreceptors have died, and cones that do not show age related loss have significantly reduced ability (7,8). Mitochondrial function in the ageing primate retina declines significantly with over a 70% reduction in adenosine triphosphate (ATP) production, which is a key energy source needed for metabolic function. This is in spite of a significant increase in glycolysis (9). These factors likely have an impact on aged human vision as improving aged mitochondrial function is associated with improved colour contrast (10) and scotopic thresholds (11).

Studies that examine patterns of retinal ageing have commonly used rodents. However, vision is not their primary sensory modality, and the animals prefer darkness. They have not evolved a high degree of central visual acuity mediated by a region of high cell density that in humans can be a focus of aged disease. But importantly, murine patterns of outer retinal cell loss are profoundly different from those in primates. In mice, unlike primates, cone photoreceptor loss is not only an ageing feature, but is initiated before rod loss(12), and rodent age related patterns of cytokine expression are completely different from that found in primates in both the retina and serum (9).

For these reasons it is important that we examine retinal ageing in the closest living relative to humans, old world primates, and that this should have a focus on mitochondria that are critical in the ageing process(1) and take account of central and peripheral region, as there is evidence for preferential and progressive accumulation of mitochondrial DNA (mtDNA) mutations in the central human retina compared to the periphery(13). Here we examine mitochondrial decline in the old world primate(*Macaca fascicularis*), describing key features and also asking if similar processes are found in the central retina, which mediates the majority of our visual function as in the periphery. We also describe key aged changes in cone photoreceptors that do not die with age and on which, because of artificial lighting, we have become increasingly dependent.

## Materials and Methods

### Animals

Primate ocular tissues were acquired from a large, long established colony of *Macaca fascicularis* maintained by UK Public Health laboratory regulated under local and U.K. Home Office regulation. The primary purpose of animal usage was different from the aims of this study and eyes were only retrieved for this study after death. The primates are of Mauritian origin. The Mauritian primates have a much smaller gene pool than those from Indonesia and age at a more rapid rate. Their normal maximum lifespan in this colony is around 18 years compared to approximately up to a maximum of 30 years in a similar Indonesian population. However, lifespan in the wild varies considerably across subspecies and in relation to habitat. However, his makes the Mauritian animals used here ideal and unique model for old world primate ageing.

Eye retrieval followed termination after sedation with ketamine (200 mg/kg) and overdose of intravenous sodium pentobarbital (50 mg/kg). The eyes were removed rapidly. One eye from each animal was rapidly dissected removing retina, retinal pigment epithelial (RPE) and choroid from the macular and defined region of the periphery. At each location, the retina was separated from RPE and choroid. Samples were taken from the central macular just beyond the fovea by approximately 2mm. Those from the periphery were outside the macular area with tissue pooled from the nasal, ventral and dorsal regions approximately4mm distant from the optic nerve head. These samples were snap frozen and stored at −80°C for different assays. The other eye was fixed in 4% paraformaldehyde in PBS. Samples were taken from 10 young animals (average age 4 years. SD ± 4 months) and 10 older primates (average age 13 years and 10 months. SD ±4 months) in this study.

### Mitochondrial extraction and isolation

Mitochondria from 5 old and 5 young adults from the central macular and peripheral regions were homogenised in pre-chilled mitochondrial storage buffer consisting of 230mM Mannitol, 70mM sucrose, 1mM EGTA, 10mM HEPES and protease inhibitor cocktail (Sigma-Aldrich, UK). The homogenate was centrifuged at 800 X g for 30 mins at 4°C and then the supernatant was centrifuged at 11,000 X g for 10 mins to precipitate the mitochondria. Mitochondrial pellet was resuspended in fresh mitochondrial storage buffer and protein concentration of each sample was measured using the BCA assay (ThermoFisher Scientific, UK).

### Mitochondrial membrane potentials

The membrane potential of isolated mitochondria from above was investigated using a coloured dye described by Figueira et al.(14)in which they follow changes in fluorescence of the membrane potential probe, Safranin O with a spectrofluorometer. Safranin O undergoes an optical shift upon its potential-dependent distribution between the external medium and intramitochondrial compartment. 20μg of mitochondrial protein was added to an incubation medium containing 220mM Mannitol, 100mM Sucrose, 1mM EGTA, 4mM K_2_PO_4_, 0.2mg/ml BSA, 20mM HEPES, 10μM Safranin O and 15mM of D-glyceraldehyde-3-phosphate, pH 7.4 and the changes were read at excitation wavelength 495nm and emission wavelength 586nm for 1 minute.

### Mitochondrial membrane permeability and light scattering

Isolated mitochondrial suspensions from above containing about 50μg protein/ml were subjected to swelling using a hypotonic solution to examine their membrane permeability. The hypotonic solution consisting of 90mM KCl, 20mM MOPS-KOH, pH 7.2 and the absorbance changes were measured spectrophotometrically at 540nm for 1 minute. The absorption rate (ΔA) and the initial rate (1 min) of mitochondrial swelling (ΔA/min) were recorded. Light scattering measurements of the mitochondria were determined from absorption values prior to the addition of the hypotonic solution.

### Immunohistochemistry

Primate eyes of 5 older and 5 young were fixed in 4% paraformaldehyde in PBS for approximately 24h. Eyes were then washed in 1XPBS, dissected and the anterior tissues removed. A horizontal strip containing the retina, RPE and choroid was removed running from the temporal periphery, through the macular and fovea into the nasal retina. These were cryoprotected in 30% sucrose and then embedded in OCT (Optimal Cutting Temperature Tissue-Tek compound, Leica). Frozen sections were cryosectioned at 10 μm and processed for immunohistochemistry using the streptavidin horseradish peroxidase complex. The sections of the eye were incubated for 1 hour in a 5% Normal Donkey serum in 0.3% Triton X-100 in PBS, pH 7.4, followed by an overnight incubation with primary antibodies in the table 1 below which was made in 1% Normal Donkey Serum in 0.3% Triton X-100 in PBS. After the primary antibody incubation, the sections were washed three times in 0.1 M PBS and then treated with 0.3% hydrogen peroxide in PBS to quench endogenous peroxidase activity. After several washes, the tissues were incubated with the appropriate biotin–SP conjugated secondary antibody(Jackson ImmunoResearch Laboratories, 1:1000) which were made up in 2% Normal Donkey Serum in 0.3% Triton X-100 in PBS, were added to the sections and incubated for 1 hour at room temperature. Negative controls were undertaken omitting the primary antibody. After the secondary antibody incubation, sections were washed several times and incubated in a ready to use horseradish peroxidise Streptavidin solution (VECTOR Laboratories, UK) for 30 minutes, followed by a peroxidase substrate solution, 3,3-diaminobenzidine (DAB) for 1 minute. Slides were mounted in glycerol and coverslipped after several washes in PBS and TBS. Sections were viewed and images captured using an Epi-fluorescence bright-field microscope (Olympus BX50F4, Olympus, Japan), where data were captured as 24-bit colour images at 3840×3072 pixel resolution using Nikon DXM1200 (Nikon, Tokyo, Japan).

**Table 1.** Antibodies used, concentration and source.

| Primary antibodies used | Concentration |  | Company |
| --- | --- | --- | --- |
|  | Western blot | Immunohistochemistry |  |
| Mouse monoclonal Tom20 | 1/300 |  | Santa Cruz Biotechnology Inc. |
| Rabbit polyclonal to VDAC | 1/2000 |  | Novus Biologicals |
| Mouse monoclonal to β-actin | 1/10000 | | Proteintech |
| Mouse monoclonal to Succinate Dehydrogenase (Complex II) | 1/500 | 1/200 | Abcam |
| Goat polyclonal to cytochrome C oxidase subunit III (Complex III) | 1/1000 | 1/500 | Santa Cruz Biotechnology Inc. |
| Mouse monoclonal to COXIV (Complex IV) | 1/1000 | 1/100 | Abcam |
| Mouse monoclonal to α-tubulin | 1/5000 | | Sigma-Aldrich |
| Rabbit polyclonal to GAPDH | 1/1000 |  | Milipore |

### Western blotting

Snap frozen retinal samples from 5 older and 5 young primates from both central macular and peripheral regions were homogenized in 2% sodium dodecyl sulfate (SDS, Sigma Aldrich, UK) with protease inhibitor cocktail (Sigma Aldrich, UK), and centrifuged at 13,000 X g. The supernatant was transferred to a new microcentrifuge tube and for Western blots of the proteins in table 1; translocase of outer membrane 20 (Tom20), a marker of mitochondria and the mitochondrial porins voltage-dependent anion channel (VDAC), proteins of mitochondrial function: complex II, III and IV. The protein concentration of each sample was measured using the BCA Protein Assay (ThermoFischer Scientific, UK). Equal amounts of protein (50ug/ml) were separated by a 10% SDS gel electrophoresis and electrophoretically transferred onto nylon membranes. Then the nylon membranes were pre-treated with 5% non-fat dried milk in 1M phosphate buffered saline (PBS), pH7.4 for 1 h and incubated overnight at room temperature with the primary antibodies in the table below followed by several washes in 0.05% Tween-20 in 1M PBS. The membranes were then incubated with the respective secondary antibodies; goat anti-mouse HRP conjugated (1/5000, ThermoFisher Scientific UK), rabbit anti goat HRP conjugated HRP (1/5000, Dako UK) and goat anti-rabbit HRP conjugated (1/5000, Dako UK) for 1 h. Immunoreactivities were visualised by exposing x-ray films to blots incubated with enhanced chemiluminescence (ECL) reagent (SuperSignal West Dura, ThermoFisher Scientific UK). Total protein profile was determined by staining blots with Ponceau S solution to check the transfer efficiency and quantification. Protein bands were then photographed and scanned. After each protein detected, the nylon membranes were then stripped with a stripping buffer containing 6M guanidine hydrochloride, 0.2% Nonidet P-40, 0.1M β-mercaptoethanol and 20mM Tris-HCL and washed several times with the washing buffer made up of 0.14M NaCl, 10mM Tris-HCl and 0.05% Nonidet P-40 before being reprobed by another primary antibody of interest (table 1).The absolute intensity of each band was measured using Adobe Photoshop CS5 extended.

### Resin embedded histology

Primate eyes of 5 older and 5 young were fixed in 4% paraformaldehyde in PBS, and then transferred to 0.1M PBS for dissection. Horizontal strip of the neural retina together with the RPE/choroidal tissues were dissected from the periphery to the macular and were processed for resin embedded plastic. They were post-fixed with 2% paraformaldehyde and 2% glutaraldehyde in PBS for 24h, followed by repeated PBS washing and dehydrated through a series of ethanol. The samples were infiltrated, polymerised and embedded in Technovit 7100 historesin (Taab Laboratories equipment, UK). Resin sections were cut at 2.5μm using a microtome, histologically stained with Acid Fuchsin (mitochondrial stain) and mounted in Depex, cover slipped.

## Results

### Complex activities

Complex activities were examined by western blot and immunostaining for complexes II, III and IV. Patterns in immunostaining for the complexes were similar across retinal layers. As expected, the most prominent feature of the immunostaining was the heavy label in photoreceptor inner segments. Staining was also present in the inner retina but largely absent from the outer nuclear layer. There were no obvious differences with age or location in the extent of the label (Figures 1–3).

**Figure 1.**
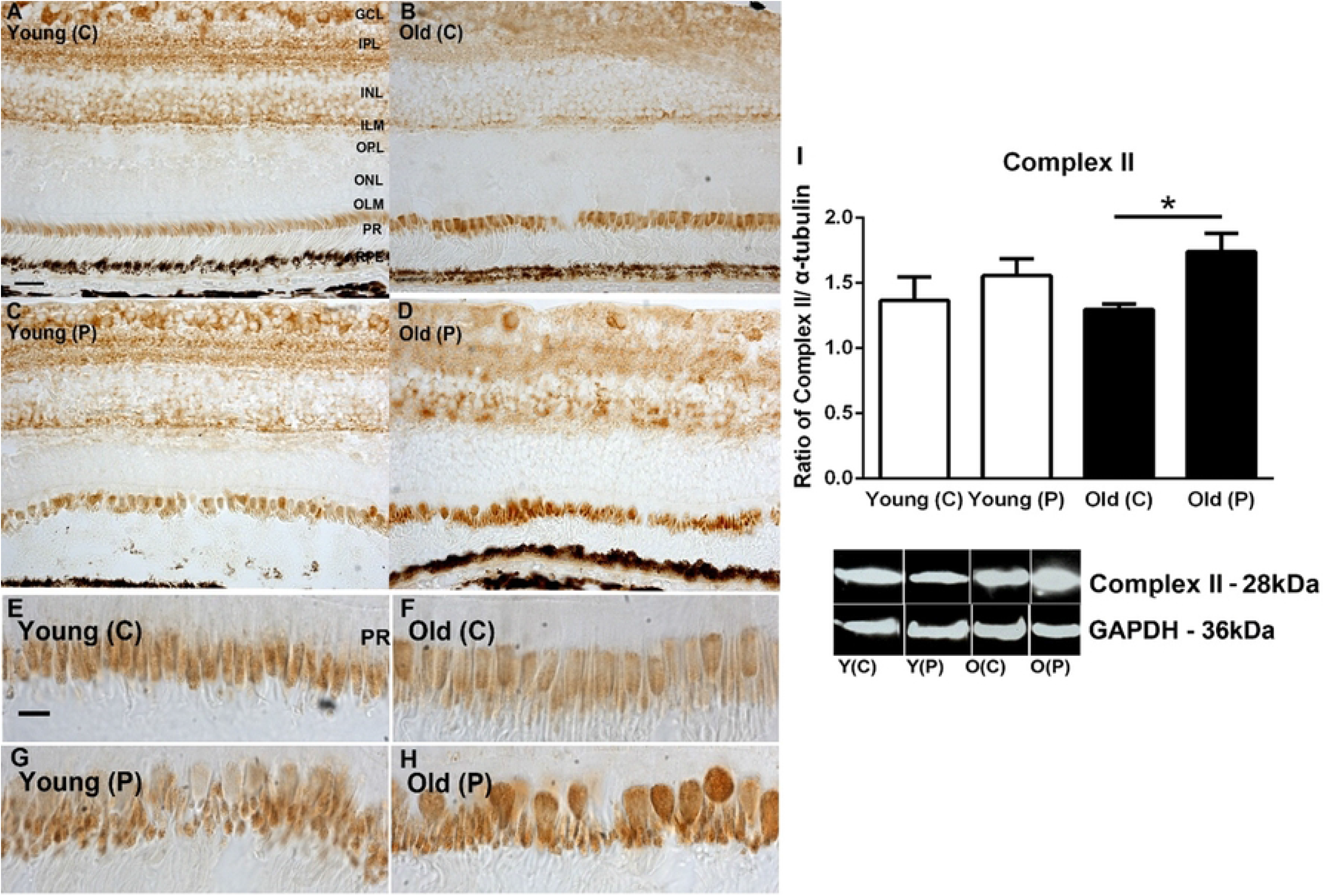
**A-D.** Immunohistochemistry of succinate dehydrogenase (complex II) activities on young and old primate central (C) and peripheral (P) retina. Staining is present mostly in the ganglion cell layer (GCL), Inner plexiform layer (IPL) and strongly in inner segment of the photoreceptor layer (PR). **E-H.** Inner segments at higher magnification. Staining appears stronger in peripheral sections particularly in the older animals. **I.** Western blot analysis for complex II confirms the immune staining with higher levels of complex II found in the periphery particularly in older animals. However, there is no evidence of age related changes. N=5 per group. Abbreviations: GCL, ganglion cell layer. IPL, inner plexiform layer. INL, inner nuclear layer. ILM, inner limiting membrane. OPL, outer plexiform layer. ONL, outer nuclear layer. ILM, outer limiting membrane. Photoreceptors, PR. Young centre, Y(C). Young periphery Y(P). Old centre, O(C). Old periphery, O (P). Statistical significance, * p<0.05.

**Figure 2.**
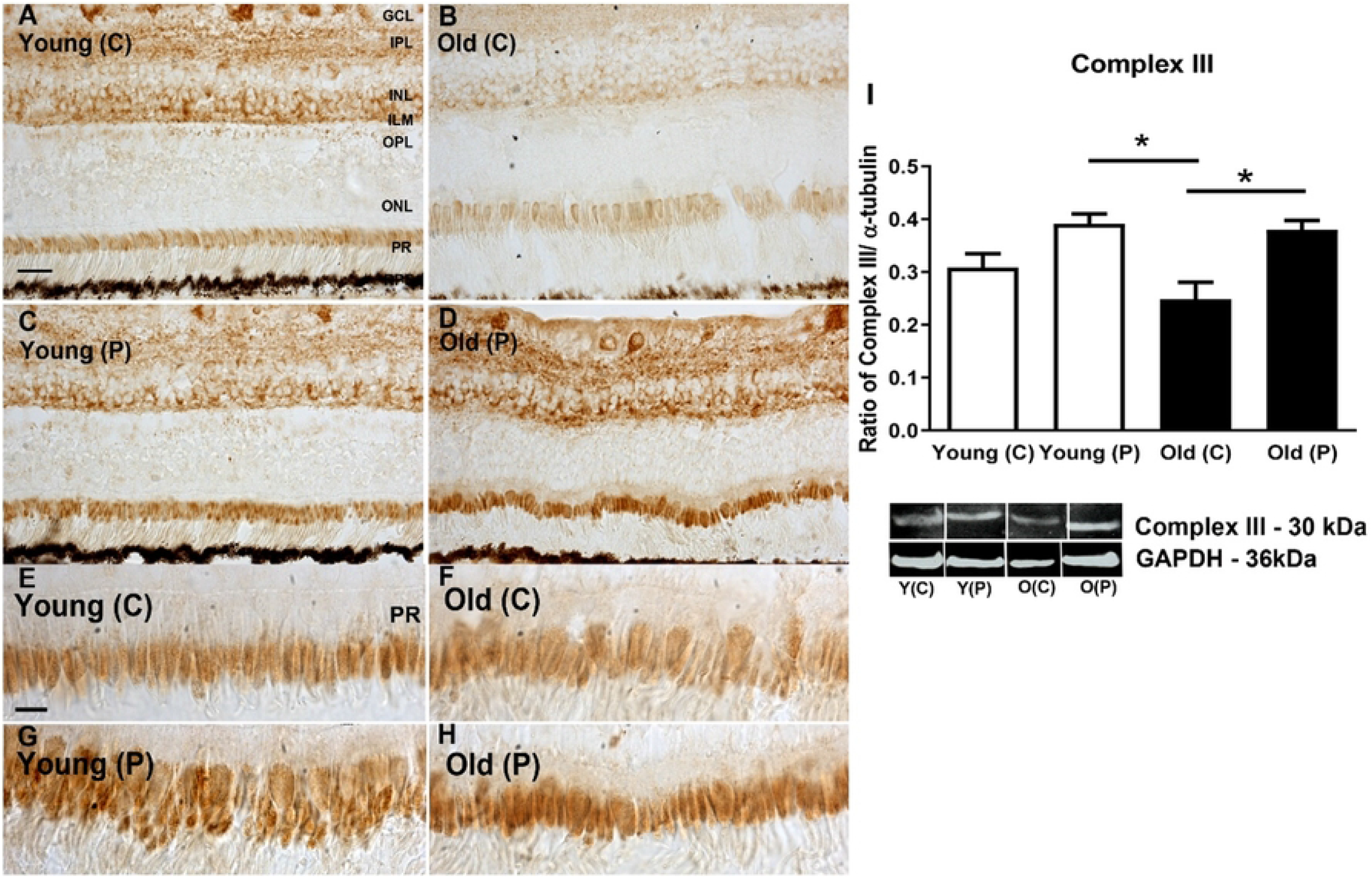
**A-D.** Immunohistochemistry of cytochrome C oxidase subunit III in young and old primate central (C) and peripheral (P) retina. Positive staining is present in the ganglion cell layer (GCL), Inner plexiform layer (IPL), inner nuclear layer (INL) and inner segment of the photoreceptor layer (PR). In both age groups, staining in the periphery appeared heavier than in the central retina. **E-H**. Images of photoreceptor inner segments at higher magnification. The peripheral images in both aged groups appeared to show heavier staining than central region. **I.** Western blot analysis confirming higher levels of expression of Complex III in the periphery of both age groups, but also revealing no age related changes in expression. N=5 per group. Abbreviations: GCL, ganglion cell layer. IPL, inner plexiform layer. INL, inner nuclear layer. ILM, inner limiting membrane. OPL, outer plexiform layer. ONL, outer nuclear layer. ILM, outer limiting membrane. Photoreceptors, PR. Young centre, Y(C). Young periphery Y(P). Old centre, O(C). Old periphery, O(P). Statistical significance, * p<0.05.

**Figure 3.**
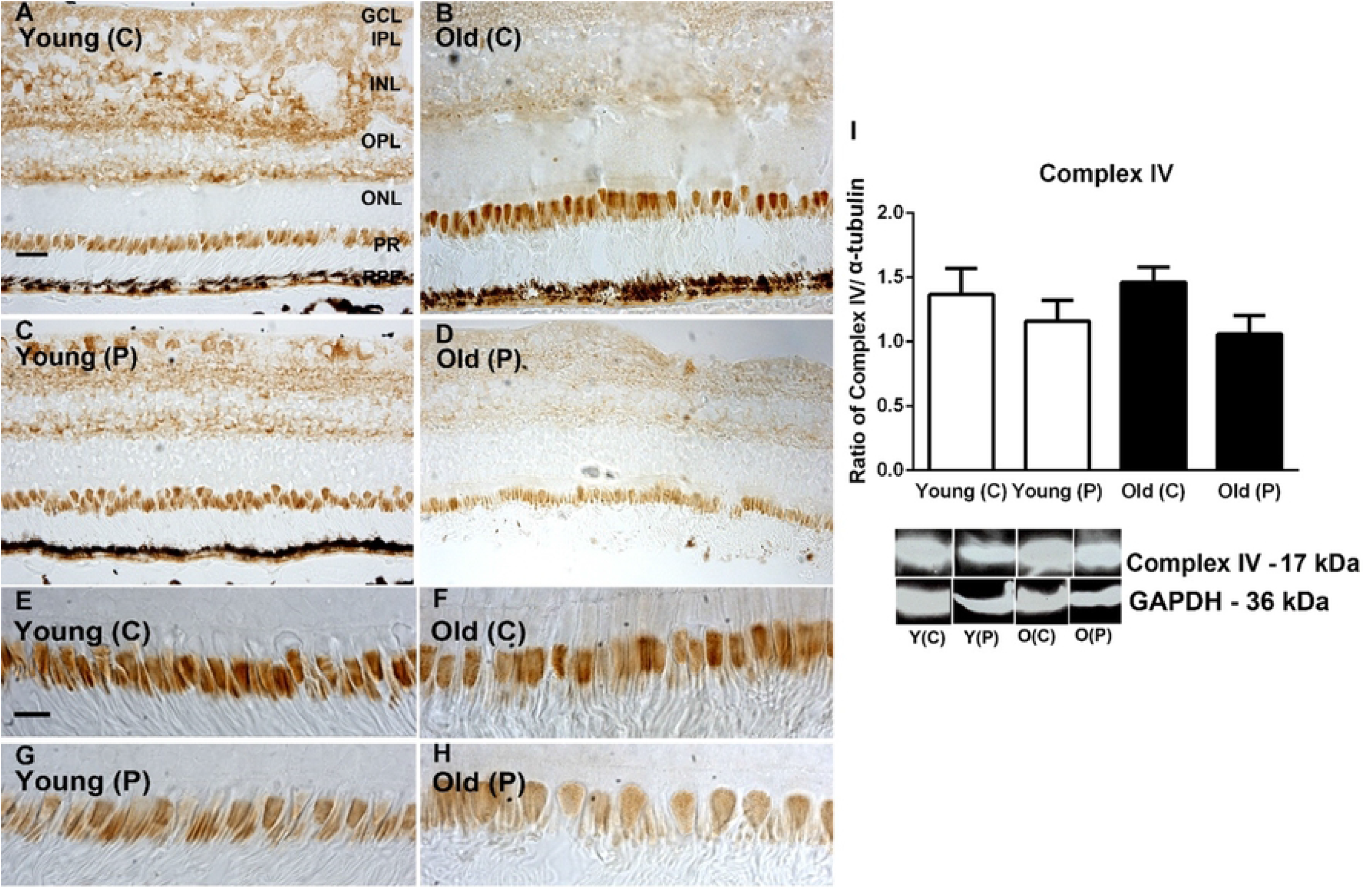
**A-D.** Immunohistochemistry of cytochrome C oxidase subunit IV in young and old primate central (C) and peripheral (P) retina. Staining is present mostly in the ganglion cell layer (GCL), Inner plexiform layer (IPL) and outer plexiform layer (OPL, also in inner segments of the photoreceptor layer (PR). Central expression appeared heavier than that in the periphery. **E-H**. Images of inner segment of photoreceptors at higher magnification. Central inner segments of both age groups appear to show higher levels of expression than those in the periphery. **I.** Immunostaining results were confirmed with Western blot showing higher levels of complex IV in central regions. However, there were no age related differences. N=5 per group. Abbreviations: GCL, ganglion cell layer. IPL, inner plexiform layer. INL, inner nuclear layer. ILM, inner limiting membrane. OPL, outer plexiform layer. ONL, outer nuclear layer. ILM, outer limiting membrane. Photoreceptors, PR. Young centre, Y(C). Young periphery Y(P). Old centre, O(C). Old periphery, O(P).

A similar analysis was undertaken with Western blot to quantify complex protein expression. Complexes II and III were consistently more abundant in the periphery than the centre in both young and older animals. In complex IV greater concentrations were found in the centre than the periphery in both young and older, but these differences were not statistically significant. These data are similar to those presented previously by us(9) where we measured the enzyme kinetic of complex IV activities, however previously the difference between young and older in the periphery did just reach statistical significance. The overall result for complex measurements was that there were no significant changes with age in spite of the 70% reduction in ATP documented in the same population of primates (Figures 1–3).

### Mitochondrial membrane potential, permeability and size

A measure of mitochondrial viability can be obtained by examining membrane potential, membrane permeability and mitochondrial size that were derived from isolated mitochondrial preparations. Generally, with age there are reductions in mitochondrial membrane potential, an increase in their membrane permeability and an overall increase in mitochondrial size. These features were seen here in the primate retinal tissues.

Membrane potentials were significantly lower in older than younger animals although there were not significant regional differences. Overall reductions across age were approximately 30-40% (Figure 4A). Reduced membrane potentials were associated with increased membrane permeability. Again, there were no significant regional differences within either age group but there was a significant increase in permeability with age of approximately 30-40% (Figure 4B).

**Figure 4.**
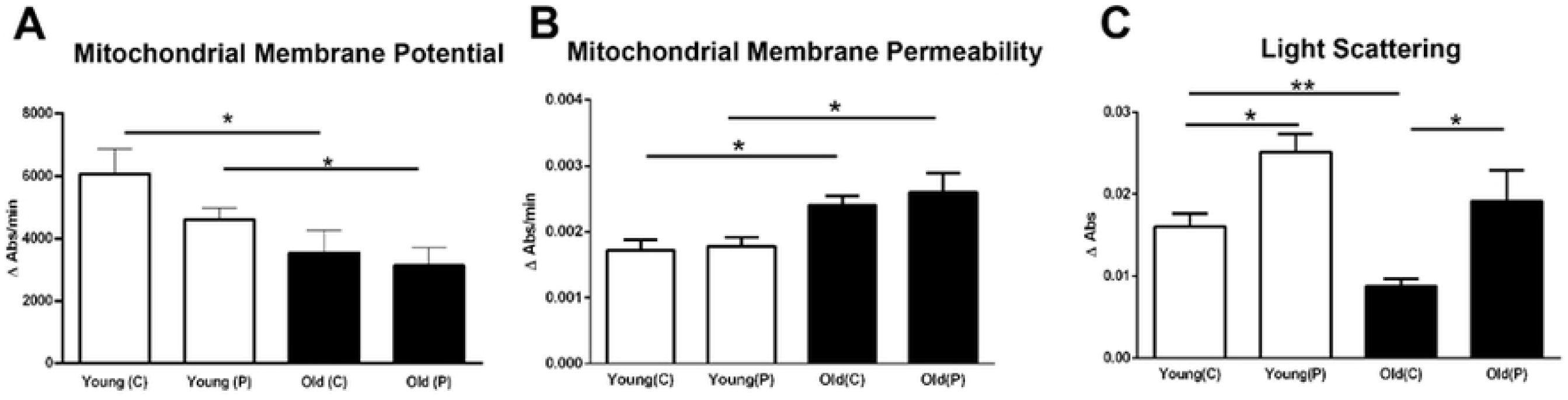
Three mitochondrial metrics were assessed from the same retinal samples across age and location **A.** Retinal mitochondrial membrane potentials from young and old primates from central and peripheral regions. An overall significant age-related decrease of between 30-40% was revealed in membrane potentials for both central and peripheral regions. **B.** Mitochondrial membrane permeability was assessed revealing an inverse relationship with mitochondrial membrane potential with permeability increasing with age across the retina associated with declining potentials. Increased aged permeability was approximately 30-40% independent of location. **C.** Mitochondrial size was determined via light scatter. Larger mitochondria scatter less light due to increased absorbance by their larger surface area. In both age groups, mitochondria were larger in the periphery than the centre. However, with age mitochondrial increased in size, however, this only reached statistical significance in the central retina. Hence, with age mitochondria have reduced membrane potential, increased permeability and in the centre increased size. N = 5 per groups. Abbreviations: Central, C. Peripheral, P. Statistical significance, * p<0.05*,** p<0.01.

Reductions in light scatter are associated with an increase in mitochondrial size. In both young and old primates, mitochondrial size was greater in the centre than the periphery and in both regions, there was an increase in mitochondrial size with age, but this was only statistically significant for the central retina (Figure 4C).

### Mitochondrial number and voltage-dependent anion channels (VDAC)

Estimating mitochondrial number in whole retinae is challenging and not realistically possible in electron microscopy (EM) or using Mito trackers that require fresh tissue. Hence, here we have used Tom20 a mitochondrial membrane marker to estimate their number. Western blot analysis shows a significant approximate 50% reduction in Tom20 between young and older retinae with no regional difference between centre and periphery in either age group, consistent with there being significant mitochondrial decline with age (Figure 5A). VDAC is a general diffusion pore for small hydrophilic molecules that opens when mitochondrial membrane potentials decline and plays a role in metabolite transport and can release cytochrome c that triggers apoptosis. When this is normalised against beta actin less is found in the periphery in young and older, and overall less in older animals (Figure 5B). But when it is normalised against Tom20 there is a significant increase with age across both retinal regions (Figure 5C). This is consistent with the notion that there is an increase in VDAC pore in age that could be indicative of increased apoptosis.

**Figure 5.**
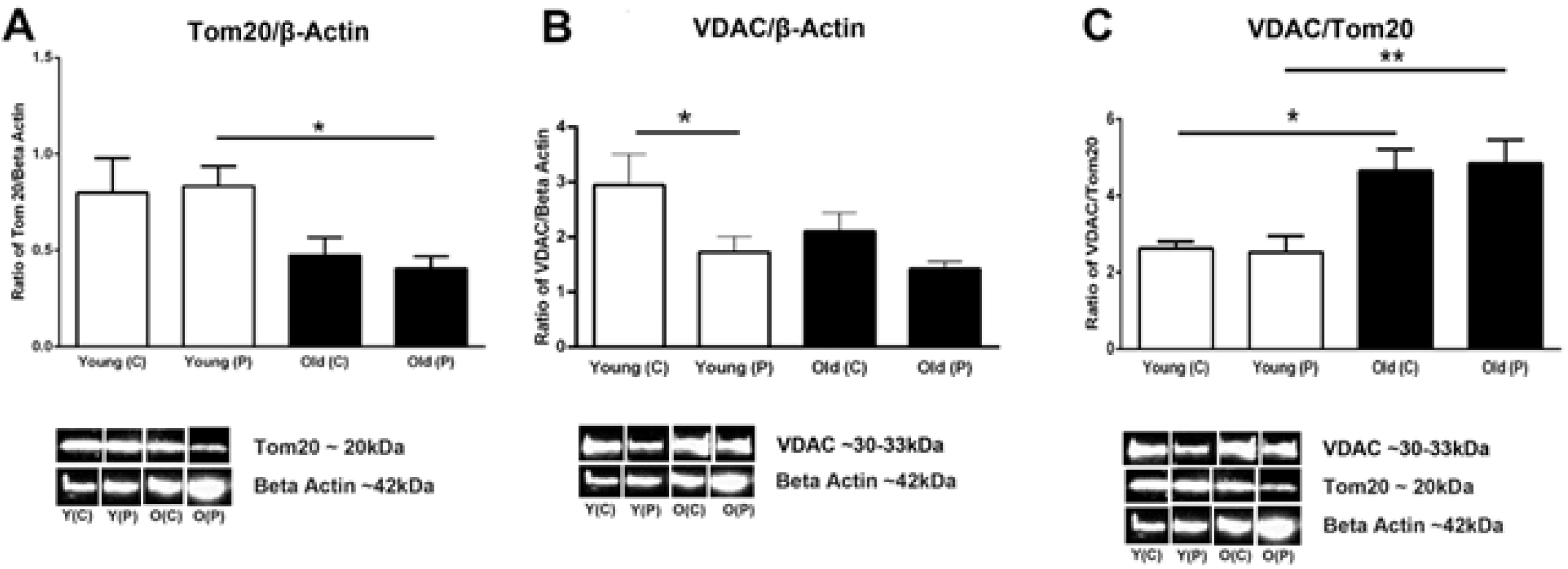
**A**. Quantitative assessment of the mitochondrial marker Tom20 normalised with β-actin using western blots. Tom20 is localised in the outer mitochondrial membrane and it recognises precursors to facilitate protein across the outer mitochondrial membrane. The graph shows that young primates have more mitochondria than the old primates do. Peripheral retina in young have significantly higher mitochondria than peripheral retina of older primates (P = 0.0143). **B.** Western blot analysis of the protein voltage-dependent anion channel (VDAC) which is localised in the outer membrane of the mitochondria and it controls hydrophilic solutes between the mitochondria and the other cell compartments. The overall results show that younger primates have more VDAC than older primates when normalised with β-actin but central retina of both young and old primates have more VDAC than the peripheral retina. The central retina of young primates have significantly more VDAC than the young peripheral retina (P = 0.0278). There was no statistical significant decrease between old primate peripheral retina compared to the centre but there is a trend that central retina has less VDAC than peripheral when normalised with beta-Actin. **C.** Western blot results of VDAC normalised with the amount of mitochondria in retina of young and old primates. It shows that older primates both in the central and periphery have significantly more VDAC/Tom20 ratio than young primates do (P = 0.0179 and P= 0.0079 respectively). N = 5 in each groups * p<0.05, ** p<0.01, *** p<0.001; C-central and P – peripheral; Y–young and O-old.

### Outer retinal structure

Mitochondria are tightly packed in the apical region of photoreceptor inner segments(15) and their density here accounts for a significant proportion of those found in the retina. With age there is significant rod loss in the primate but there is no evidence for cone loss (7,16), although cone function declines (17). Hence, we ask what mitochondrial features of ageing are present in cones that may explain their loss of function.

A key feature of the ageing outer retina in the primate was the loss of content of the proximal inner segment in many cone photoreceptors. This region is normally occupied by the endoplasmic reticulum (15) that is critical for the maintenance of fission (OPA1) and fusion (Fis1) processes in mitochondrial dynamics. This feature of the aged inner segment was twice as prevalent in the peripheral compared to the centre (Figure 6).

**Figure 6.**
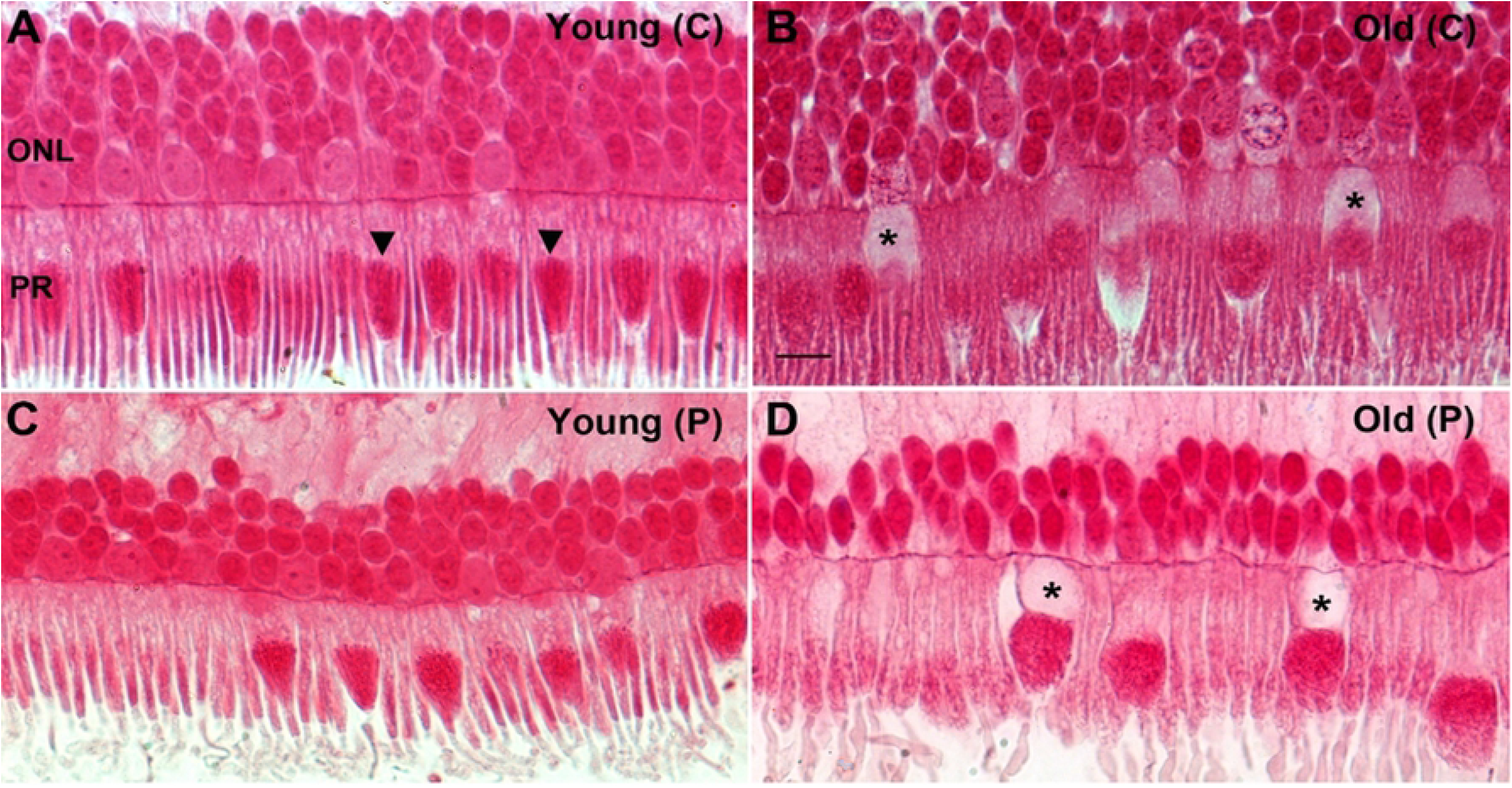
Representative photomicrographs showing age related changes in primate retina. Retina from young and old primates were processed and embedded in plastic, sectioned and stained with Acid Fuchsin (mitochondrial stain). **A.** Photomicrograph of a young primate taken at the centre of the retina showing compact and elongated photoreceptors. The mitochondria (arrowheads) are thin, tubular and elongated. **B.** Photomicrograph from old primate central retina showing the photoreceptors are swollen and some of them there seem to have less Golgi and endoplasmic reticulum in the inner segment (* asterix). The mitochondria are sparse, fragmented and rounder in shape. **C.** Photomicrograph of young primate retina from the peripheral. The mitochondria have thin tubular and elongated shape. **D.** Photomicrograph of old retinae from the peripheral area. The photoreceptors are swollen and some have a void in them at the Golgi and endoplasmic reticulum area of the inner segments (* asterix). The mitochondria appear fragmented and rounder in shape.

It has been noted that in human retinae some cones have a displacement of their nuclei through the outer limiting membrane(18,19). Figure 7A shows a series of adjacent cones that display progressive translocations of their nuclei through the outer limiting membrane into the inner segment. Star (*) in this figure is a rod that also appears to have a nucleus breaking through the outer limiting membrane. In Figure 7B, a cone nucleus has fully transitioned into the inner segment. In Figure 7C, the cone indicated by the arrow has not only transitioned into the inner segment but is directly adjacent to the mitochondrial population and the vacuous region is present in the outer nuclear layer. On the right had side other vacuous regions can be seen crossing into the outer nuclear layer.

**Figure 7.**
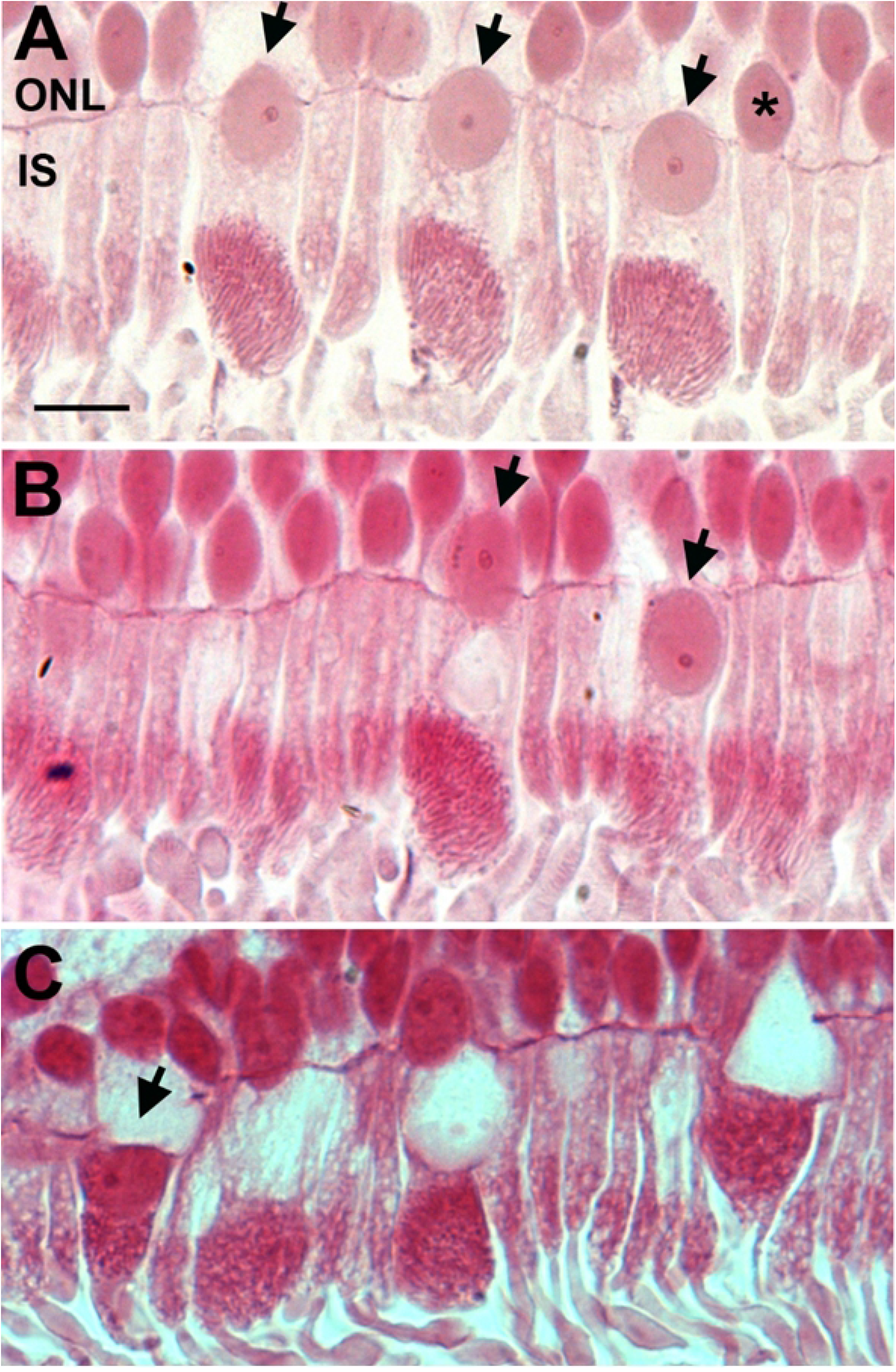
Displacement of nuclei from the outer nuclear layer to the inner segment. Representative photomicrographs of old primate peripheral retina showing displacement of the nuclei from the outer nuclear layer (ONL) to the inner segment (IS) of the photoreceptors (black arrows). The retina were sectioned at 2.5 μm and stained with Acid Fuchsin. **A**. Nuclei displacements seem to occur mainly in cones (black arrows) and rarely in rods (*asterix). It is observed that nuclei displacement occur more in the old primate periphery than in young peripheral retina. **B**. Micrograph shows that the nucleus has completely crossed the outer limiting membrane into the inner segment, in the area of endoplasmic reticulum and Golgi (the black arrow on the right). **C** shows that the nuclei has migrated and is sitting on top of the mitochondria (black arrow). Scale bar = 10μm.

## Discussion

The results of this study show that in spite of a 70% reduction in retinal ATP in the same colony of old world primates as we have used here(9), complex activity is not reduced. However, other mitochondrial changes are marked with age with reduced membrane potential and increased mitochondrial membrane permeability, along with evidence of mitochondrial swelling as shown by reduced light scatter. Further, aged changes in Tom20 are consistent with an aged decline in mitochondrial number, and when mapped against VDAC the data are consistent with opening of membrane channels that can be associated with apoptosis. This study also shows significant structural changes in regions that would normally be occupied by endoplasmic reticulum in cones. Here the nucleus in some cells crosses the outer retinal margin and enters the inner segments and can become embed in the mitochondrial population. The endoplasmic reticulum is a key player in regulating mitochondrial dynamics and these structural changes may undermine this process.

As primate cones do not die with age, but photopic function declines significantly (7,16,17), it is likely that the mitochondrial decline documented here, along with structural changes in the inner segment are associated with visual decline. The structural changes in the inner segment appear to be unique to primates, and have not to our knowledge, been reported in other animals. This reinforces the need to examine retinal ageing in a model appropriate for humans.

Our primates were of Mauritian origin and most likely have a restricted gene pool because of a higher rate of inbreeding within and closed island population. While the Indonesian macaques have longer life spans with a maximum around 30 years the average age of death in our long established population is close to 18 years. By this time the primates display a wide range of hallmarks of ageing including reduced fertility, grey hair, arthritic joints and an advanced browning of the lens. As such they are ideal models for ageing research.

A challenging question arising from the data in this study is the apparent discrepancy between the 70% reduction in retinal ATP found with age by us in a previous study on animals from the same colony as used here(9), compared to the failure to find any significant age related reduction in mitochondrial complex activity. Here there are a number of resolving possibilities. First, it is possible that there are the same number of electron transport chains, but fewer ATPase molecules to convert energy held within H ions, into energy trapped within ATP. Second, ATPase pumps work both ways, as such there could be a continual breakdown of ATP alongside production. Third, a component of the electron transport chain could be lost with age, as such even though most proteins are there, the chain is disrupted. At this stage it is unclear which of these mechanisms, or any alternative, is responsible for the discrepancy between ATP production and complex activity.

In our previous study (9), we showed that there is a significant drop in the enzyme activities of complex IV with age. Here the results showed that the same amount of enzyme complex IV is present in both young and older primates. This indicate that even the enzymes are present in the same amount, but their activity is significantly reduced. There is a further complication. It is important that the data presented here for mitochondrial function is seen as a snapshot of aged differences. Mitochondria change their function over the day and this includes complex activity, ATP production and ADP/ATP ratios(20). The primates in this study were all culled between 9am and 10.30am. However, sacrifice at different times may result in different patterns between the age groups, as we have little idea how the mitochondrial metrics in the primate retina change over the day or how any daily patterns in these in young animals are impacted by age.

The reductions in mitochondrial number, reduced membrane potential, increased membrane permeability and mitochondrial size are all consistent with age related changes (21–23). These metrics are also consistent with the increase in VDAC expression. Taken together they demonstrate declines in mitochondria with age. However, the one striking factor is that this study offers almost no evidence for the central retina suffering a greater degree of age related decline than found in the periphery as has been proposed (24). But there are a number of differences between the Barron et al study finding regional differences and our own. Barron et al examined the fovea directly, which we did not, our analysis was confined to the central macular. Further, their analysis was of mtDNA deletions and cytochrome c oxidase deficiently. We made no attempt to examine mtDNA deletions, although our inability to find any significant reductions in cytochrome c oxidase (complex IV) is perplexing. But there are a number of different factors. Barron et al. studied human postmortem human tissue spanning ages, collected a number of hours after death, and likely at different times of the day, while our study was taken at the point of death from animals in only two defined categories of age. Being human tissue, it would have suffered from varied environmental factors over life such as diet, different patterns of exercise and potentially smoking. Our primates experienced almost no environmental variability and were maintained on a standard vegetable-based diet over life. Taken together, this array of differences makes the two studies hard to compare.

Overall, the symmetry of changes between the two retinal regions used in this study in the analysis of mitochondrial extracts is marked in each of the data sets we present with the exception of changes in mitochondrial size. Here there was evidence of an increase in size in central mitochondria compared to the periphery. Such changes are associated with pathology(13,23). But this was the only instance of differential mitochondrial behaviour between the two regions examined.

Interactions between the endoplasmic reticulum and mitochondria are key in regulating mitochondrial biogenesis (25,26) and their disruption may lead to neurodegeneration and aged disease (27–29). The endoplasmic reticulum is located between the nucleus and the mitochondrial population in the inner segment, although elements of the endoplasmic reticulum project up through mitochondrial population, particularly those located proximally to the nucleus(15). The apparent loss of the endoplasmic reticulum in many cones, particularly those in the periphery is likely to be associated with disruption to mitochondrial dynamics that play an important role in autophagy and the maintenance of mitochondrial health.

Apparent loss of the endoplasmic reticulum was associated with movement of the nucleus across the outer limiting membrane to fill this void. This was obvious in cones, but also present in some rods. Nuclear translation across the outer limiting membrane has been noted before in aged human retinae (18,19), and as in this study, it is primarily a feature of the peripheral retina. In some cases, the nucleus embeds in the mitochondrial population and the vacuous region is left behind and crosses the outer limiting membrane into the ONL. These structural changes have not been noted in other aged animals, particularly mice that are commonly used as aged models

There is no evidence for cone loss in the ageing primate retina including humans(7,16). Counting primate cones is often undertaken from whole mounted retinae where the large cone inner segment distinguishes them from rods (7), or they can be identified when stained for a cone opsin and the outer segments are counted (7). In neither preparation would the structural features we have described be readily identified. However, given the marked changes that occur to the internal structure of many aged cones it would be surprising if these cells functioned normally. Such changes along with those in mitochondrial integrity including reduced membrane potential and their increased permeability are likely to impact on ageing human photopic vision and may be related to aged reductions in colour sensitivity (17). An additional factor that is relevant to this study is the finding that the spatial organisation of mitochondria in mammalian cones has the ability to act as light guide, focussing the light path onto the outer segments. It is likely that the changes we describe here on aged mitochondria impact upon this ability (30).

Given that humans now have the ability to regulate environmental light 24hrs a day, we rarely use our rod population. This places emphasis on developing strategies to maintain our cone populations in societies who average age is increasing progressively.

